# BCAT1 regulates glioblastoma cell plasticity and contributes to immunosuppression

**DOI:** 10.1101/2023.04.28.538788

**Authors:** Pavle Boskovic, Nathalie Wilke, Peter Lichter, Liliana Francois, Bernhard Radlwimmer

**Author notes:** Correspondence to: Bernhard Radlwimmer Full address: Im Neuenheimer Feld 280, 69120 Heidelberg, Germany. These authors jointly supervised this work (Liliana Francois, Bernhard Radlwimmer).

## Abstract

Glioblastoma is the most common malignant brain tumor in adults. Cellular plasticity and the poorly differentiated features result in a fast relapse of the tumors following treatment. Moreover, the immunosuppressive microenvironment proved to be a major obstacle to immunotherapeutic approaches. Branched-chain amino acid transaminase 1 (BCAT1) is a metabolic enzyme that converts branched-chain amino acids into branched-chain keto acids, depleting cellular α-ketoglutarate and producing glutamate. BCAT1 is expressed in and drives the growth of glioblastoma and other cancers. Here we show that low-BCAT1 expression correlates with differentiated glioblastoma subtypes and its knockout (KO) results in a differentiated phenotype in human and mouse glioblastoma cells. Consistent with these observations, Bcat1-KO mouse glioblastoma cells were highly susceptible to serum-induced differentiation *in vitro*. The transition to a differentiated cell state was linked to the increased activity of TET demethylases and the hypomethylation and activation of neuronal differentiation genes. Orthotopic tumor injection into immunocompetent mice demonstrated that the brain microenvironment is sufficient to induce differentiation of Bcat1-KO tumors *in vivo*. In addition, the knockout of Bcat1 attenuated immunosuppression, allowing for an extensive infiltration of CD8^+^ cytotoxic T-cells and complete abrogation of tumor growth. Additional analysis in immunodeficient hosts revealed that both Bcat1-KO-induced differentiation and immunomodulation contribute to the long-term suppression of tumor growth. In summary, our study demonstrates that BCAT1 promotes glioblastoma growth by blocking tumor cell differentiation and sustaining an immunosuppressive microenvironment. These findings suggest novel modes limiting glioblastoma phenotypic plasticity and therapeutic failure through targeting BCAT1.

**Importance of the study:** High expression of BCAT1 occurs in many tumor entities and is related to aggressiveness, proliferation and invasion of tumor cells. In this study, we show that its expression is crucial for the continuous growth of glioblastoma cells by preventing their differentiation. Furthermore, we show that the expression of BCAT1 modulates the tumor immune microenvironment, suppressing the CD8 T-cell response. BCAT1 knockout causes glioblastoma cell differentiation and a persistent CD8 T-cell response, which is sufficient to abrogate tumor growth and prolong survival in *in vivo* immunocompetent and immunodeficient models, respectively. Our findings consolidate BCAT1 as a major player in glioblastoma and highlight its importance as a potential future target of research in this and other tumor entities.

**Key Points:**

- BCAT1 expression maintains poorly differentiated features of glioblastoma cells and provides resistance to differentiation.
- Expression of BCAT1 in glioblastoma cells contributes to the immunosuppressive features of the tumor.

## Introduction

Cellular plasticity and poorly differentiated cell features are key aspects of cancer cells, largely considered drivers of intratumoral heterogeneity and treatment resistance^1^. Thus, a better understanding of the molecular mechanisms underlying phenotypic plasticity in cancer is needed to identify novel opportunities of overcoming therapy resistance.

Glioblastoma is the most common primary central nervous system malignancy^2^. Despite an aggressive treatment plan consisting of maximal surgical resection, radiotherapy and chemotherapy, the average survival time of 15 months has not significantly improved since the introduction of temozolomide nearly 20 years ago^3^. Furthermore, novel therapies applying small molecule inhibitors in targeted approaches or different immunomodulatory treatments did not result in the prolonged survival of patients^4–6^. These treatment failures are attributable to a high level of glioblastoma cellular plasticity and tumor heterogeneity^7^, features well reflected in their capability to appropriate differentiated features and neuronal signaling during migration and infiltration and revert to a highly proliferative mesenchymal state once situated^8^. Moreover, the differentiation state of glioblastoma cells and their molecular subtypes have been proposed to shape the microenvironmental milieu and immunosuppression, another major obstacle in glioblastoma treatment^9, 10^. Attempts have been made to overcome cellular plasticity and induce glioblastoma cell differentiation through kinase inhibition^11^, treatment with differentiation factors^12^, or overexpression of differentiation-program regulators such as NEUROD1 and ASCL1^13, 14^. However, treatment effects were reversible upon exogenous factor withdrawal, rendering these options clinically irrelevant^12^.

The branched-chain amino acid transaminase 1 (BCAT1) is a metabolic enzyme expressed ubiquitously during embryonic development and in a tissue-restricted manner in adults, primarily the brain^15, 16^. By converting branched-chain amino acids (BCAA) into respective branched-chain ketoacids, BCAT1 depletes cellular α-ketoglutarate (αKG) and produces glutamate. Despite its limited expression in adult tissues, BCAT1 is overexpressed widely across tumor entities, including glioblastoma^17–21^. Furthermore, its overexpression has been described to promote tumor cell proliferation and invasiveness, through regulating the availability of BCAAs and consequently the mTOR pathway and autophagy, or through the glutamate-dependent promotion of GSH production^18–20, 22, 23^. In addition, we previously showed that BCAT1 maintains acute myeloid leukemia (AML) stem cells by the depletion of αKG and the consequent alteration of DNA methylation through the reduced activity of ten-eleven translocation (TET) methylcytosine dioxygenases^19^.

Here we highlight BCAT1 as a regulator of glioblastoma cell state plasticity that maintains the poorly differentiated characteristics of glioblastoma cells and contributes to the immunosuppressive tumor microenvironment. Using animal models, we show that both these mechanisms contribute significantly to adverse survival. These findings help define mechanisms by which BCAT1 drives tumor growth and suggest that targeting its function could provide an opportunity of addressing phenotypic plasticity and therapy resistance in glioblastoma.

## Materials and methods

### Cell lines

The U251-MG (U251) human glioblastoma cell line was obtained from ATCC and cultured in low-glucose DMEM (Sigma, D5921) with the addition of 10% FCS, 1x penicillin-streptomycin (P/S) and 0.5mM L-glutamine at 37°C and 10% CO_2_. The mouse mGB2 glioblastoma neurosphere cell line was cultured as described previously^24^ (DMEM/F12 (Gibco, 11330-057), 1x P/S, 1x N2 (Life Technologies, 17502048), 20ng/mL FGFb (PreproTech) and EGF (Life Technologies)). The cells were cultured at 37°C with 5% CO_2_.

### Generation of CRISPR/Cas9 knockout and shRNA knockdown cells

The generation of the BCAT1-KO U251 cell line was described previously^25^. Knockout of Bcat1 in the mouse mGB2 cell line was performed using a doxycycline-inducible CRISPR/Cas9 construct TLCv2 (addgene, 87360) with the single guide RNAs (sgRNA) 5’- CACCGGCTGACCACATGCTGACG-3’ for the control construct and 5’- CACCGGTATTACTGATATTGGTGGG -3’ for the Bcat1-KO construct. Cas9 expression was induced with doxycycline for 48h, after which single-cell clones were screened using Western blots. Cells were lysed in RIPA lysis buffer and separated on a 4-12% SDS-PAGE gel with Sample Buffer (Life Technologies, NP0007) and the reducing reagent (Life Technologies, NP0009). Proteins were transferred to a pre-activated PVDF membrane and blocked with 5% milk in TBS-T (tris-buffer saline, 0.1% Tween20). Primary antibody incubation with anti-BCAT1, anti-αTubulin and anti-V5- Tag was performed in 5% milk in TBS-T at room temperature for 4h. Secondary antibody incubation with anti-rabbit-HRP and anti-mouse-HRP was done at room temperature for 1h in 5% milk in TBS-T and the signal developed using an ECL mixture (Life Technologies, 32132). For a detailed list of antibodies see Supplementary Table 2. The Tet1 and Tet3 knockdown cell lines were produced using a constitutive pLKO.1 vector with blasticidine resistance expression (addgene, 26655) and 3 different shRNA sequences (Supplementary Table 3) per Tet protein designed using the Broad GPP web portal (https://portals.broadinstitute.org/gpp). mGB2 Bcat1-KO cells were transduced with either the shTet constructs or the control shRNA targeting Luciferase (shLuc), and the Control cells only with the shLuc construct. Cells underwent antibiotic selection 48h following transduction.

### *In vitro* cell differentiation

For the differentiation of the mGB2 cells, cells were seeded in plates or on coverslips pre-coated with ECM (Sigma-Aldrich, CC131) to promote attachment in either the standard medium containing growth factors (SC condition) or in the DMEM/F12 medium containing 5% FCS without any additional growth factors (FCS condition). All differentiation experiments were performed 8 days after the initiation of differentiation, during which time the medium was exchanged every other day with a reseeding at day 4. For recovery experiments, the FCS medium was removed, and the cells washed after 8 days of differentiation and replaced with the SC medium for the indicated time.

### RT-qPCR

Cellular RNA was isolated using the QIAGEN RNeasy Mini kit with on-column DNA digestion. 500- 1000ng of total RNA were used for cDNA synthesis (NEB, E6560S). 12.5ng of cDNA (according to RNA starting amount) per reaction was used for the qPCR reaction with the PrimaQUANT CYBR qPCR mix and 0.5µM forward and reverse primers (primer sequences can be found in Supplementary Table 1). The qPCR was performed on the QuantStudio5 and the Ct threshold for each gene was determined automatically by the QuantStudio design and analysis software. The ddCt values were calculated by normalizing to a housekeeper gene (*Tbp*) and the condition specified in the experiment. Fold change was calculated as 2^-ddCt^.

### Click-iT EdU proliferation assay

The Click-iT EdU assay for flow cytometry was performed according to manufacturer’s instructions (ThermoFisher Scientific, C10643). Briefly, cells were incubated with 10µM EdU for 1h. After the incubation, the cells were detached, fixed in 4% PFA and permeabilized in PBS-T (0.1% TritonX-100 in PBS) for 1h at room temperature. After washing with 1% BSA in PBS, cells were resuspended in the Click-iT reaction cocktail according to the specifications for the AF647-azide and incubated for 30min at room temperature. After washing, cells were processed on the BD LSRFortessa flow cytometer.

### Cell immunofluorescent stainings

For immunofluorescent stainings of cultured cells, the cells were fixed on coverslips in 4% PFA for 15min at room temperature and blocked and permeabilized using Blocking Buffer (5% BSA, 0.02% sodium azide, 0.5% TritonX-100 in PBS). Primary antibodies were diluted in 0.2X Blocking Buffer and incubated with the cells over night at 4°C in a humid chamber. Secondary fluorescently labeled antibodies were diluted in 0.2X blocking buffer and incubated with the cells for 1h at room temperature. Cover slips were mounted on slides using NucBlue-containing mounting medium (Life Technologies, P36985). Imaging was performed using the Leica SP8 confocal microscope. For a detailed list of antibodies see Supplementary Table 2.

### RNA sequencing and analysis

RNA sequencing of the U251 cells was performed using the Illumina HiSeq 2000 V4 and of the mGB2 cells with the Illumina NovaSeq 6000 SP. Library preparation and sequencing was performed by the DKFZ Genomics and Proteomics Core Facility (GPCF) using the TruSeq RNA Library prep kit. Reads were aligned by the DKFZ Omics IT and Data Management Core Facility using the One Touch Pipeline^26^.

Pre-processing, filtering, and the normalization of the data was done in R using *limma*^27^ and *DESeq2*^28^ packages. Pre-ranked GSEA analysis was performed using the *fgsea* package^29^ and the human MSigDB collections^30^. For the molecular glioblastoma subtype gene set, signatures were collected and combined from several publications^8, 31–34^.

The TCGA-GBM data was quarried for samples which included Illumina HiSeq sequencing results using R with packages *TCGAbiolinks, TCGAWorkflow* and *TCGAWorkflowData*^35^. Samples were pre-filtered according to the IDH status and downstream processing was performed as detailed above. Normalized patient-derived glioblastoma stem cell data was acquired from the HGCC database^36^.

### DNA methylation array and analysis

DNA methylation of the mGB2 cells was determined using the Infinitum Mouse Methylation BeadChip in triplicates by the GPCF. Data processing and differential methylation analysis was done in R using the *minfiData*, *missMethyl* and *DMRcate* packages adapted for the appropriate mouse array^37–39^. Pre-processing was done using the β-values and the statistical analysis and differential expression using the M-values. Gene enrichment analysis was done with the *methylGSA* package^40^.

DNA methylation of the U251 cells was assessed using the Infinitum HumanMethylation450 Bead Chip with a single sample per condition. Pre-processing was performed as detailed above.

### Nuclear αKG measurements

Nuclear aKG levels were determined using the αKG FRET-sensor TC3-R9P previously developed in our lab^41^. The SV40-derived nuclear localization signal (NLS) peptide (PKKKRKV) was inserted 3 times at the C-terminus of TC3-R9P. Control and BCAT1-KO U251 cells were reverse-transfected with 1µg/ml of TC3-R9P-3NLS pDNA using 1:2 of Trans-iT (Mirus). 48h after transfection, cells were imaged using a Leica SP8 laser scanning confocal microscope with an incubation chamber with controlled temperature set at 37°C and 5% CO_2_. Cells were excited with a UV diode (405nm) and emission was detected sequentially between frames at 450–490nm (CFP) and 520–590nm (FRET). Image processing and calculation of the FRET ratio was performed with Fiji (ImageJ). The ratio was calculated by dividing the mean intensity of the FRET signal by the CFP signal (FRET/CFP).

### 5-hmC immunofluorescence

Control and Bcat1-KO mGB2 cells were differentiated for 8 days and seeded on pre-coated cover slips. The cells were fixed with 4% PFA and permeabilized in PBS with 0.1% TritonX-100 (PBS-Tx). Antigen retrieval was performed with 2N HCl at 37°C for 30min and the acid was neutralized with 1M Tris-HCl (pH 8.3) for 10min. Blocking was performed in 5% goat serum PBS-Tx for 30min at room temperature followed by primary antibody staining in 5% goat serum PBS-Tx overnight. Secondary antibody staining was performed for 1h at room temperature in 5% goat serum PBS-Tx, and the cover slips were mounted on microscopy slides using NucBlue-containing mounting media.

### Mice

C57BL/6 and NSG female mice were acquired from Janvier labs. The mGB2 tumor transplantations were performed as previously described^24^. Briefly, mice were anesthetized with isoflurane and orthotopic transplantations were performed with 0.6*10^6^ cells in 2µl of PBS over 10min using a microsyringe at the coordinates (0,2,3) according to the bregma. *In vivo* bioluminescent measurements were performed using the IVIS system. Mice were injected intraperitoneally with 150mg/kg luciferin in PBS. Imaging was performed 10min following the injection with 3min exposure time.

All animal experiments and procedures were approved by the local authorities (Regierungspräsidium Karlsruhe, Germany) under the animal protocol G314-19.

### Mouse sample preparation and immunofluorescent staining

Upon reaching end point criteria according to the approved protocol, and before succumbing to the disease, mice were sacrificed with CO_2_ and the brain isolated and fixed in 4% methanol-free PFA for 24h at 4°C. The brains were cryoprotected in 30% sucrose and frozen in OCT on dry ice. Samples were kept at -80°C until further processing. Tissue sections were performed at -20°C with section thickness of 6-8µm and placed on SuperFrost Plus microscopy slides.

Immunofluorescent stainings were performed as described previously for cultured cells. For a detailed list of antibodies see Supplementary Table 2.

### Image processing

The majority of the image processing was performed using Fiji (ImageJ) software. Nuclear segmentation of images was performed using NucBlue-stained nuclei with the watershed segmentation method. Mean fluorescent intensity was measured in the appropriate channel per the defined Region of interest (ROI). The positivity threshold was determined based on the negative control.

QuPath software (version 0.3.2) was used for processing whole brain and tile scan immunofluorescent labeling. For quantification purposes, the Cell Detection algorithm was run using NucBlue, and cell positivity for Iba1 or CD8 was determined using the Nucleus: staining method with a consistent threshold between images.

### Statistical analysis

Data presented are shown as means with error bars indicating standard deviation or as Tukey plots with error bars indicating 1.5 IQR (interquartile range) values, unless specified otherwise in the figure legends. Multiple comparison testing was performed with one-way or two-way ANOVA analysis and pair-based comparisons were made with Tukey’s post-hoc test. Comparisons between two conditions were performed using a two-tailed unpaired student’s t-test. Significance levels are denoted with * symbols according to the figure legends.

### Data availability

The data that support the findings of this study are available from the corresponding author upon reasonable request.

## Results

### BCAT1 expression correlates with glioblastoma cell state

To better characterize the role of BCAT1 in glioblastoma, we analyzed its expression in a cohort of 141 sequenced tumors of the Cancer Genome Atlas Glioblastoma Multiforme (TCGA-GBM) dataset with available RNA-sequencing data. First, we stratified the samples based on BCAT1 expression and assigned the top and bottom 20 samples as BCAT1^high^ and BCAT1^low^, respectively (Figure 1A). Multidimensional scaling (MDS) analysis showed distinct clustering of the two groups, indicating divergent expressional patterns of the tumors based on BCAT1 expression status (Figure 1B).

**Figure 1.**
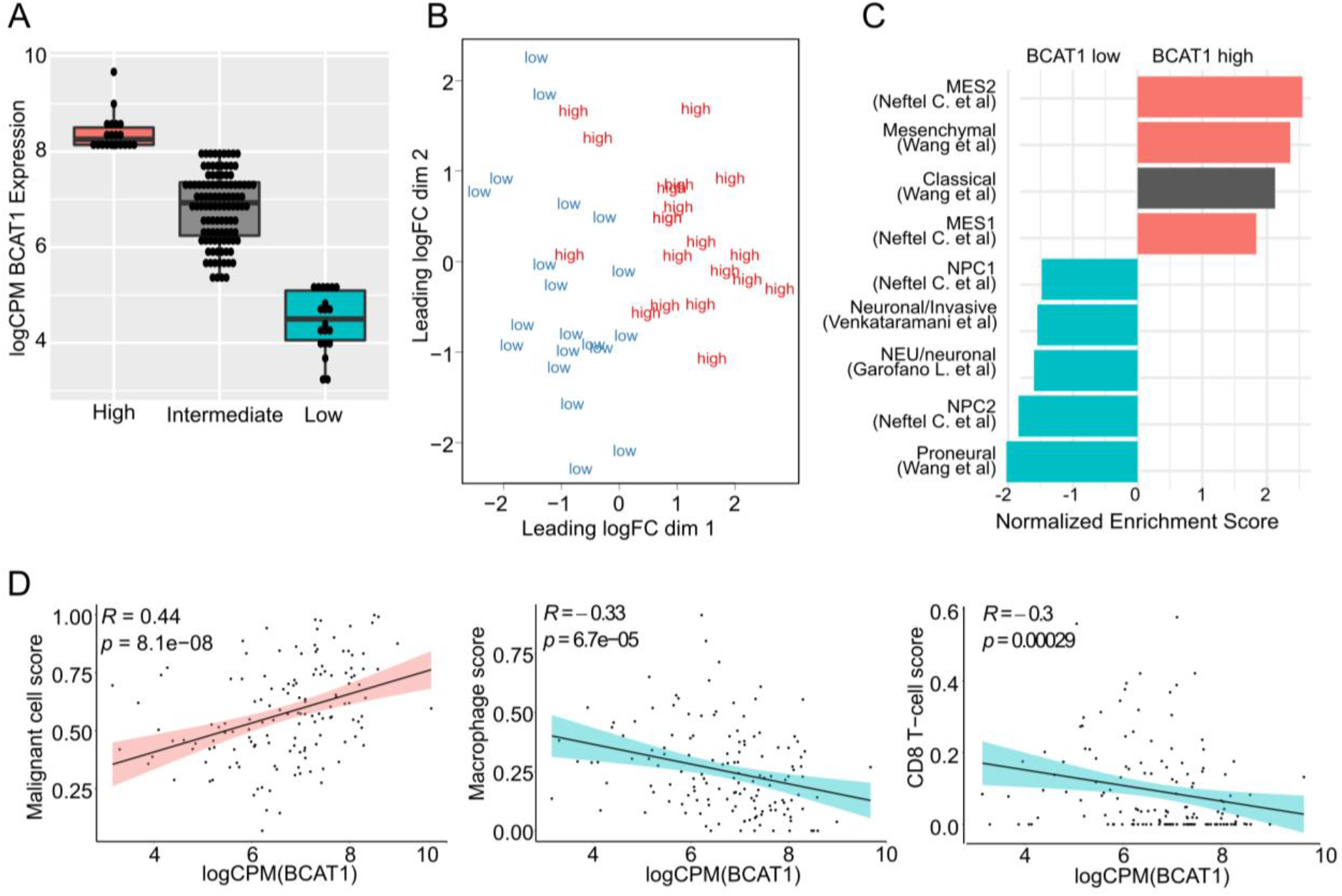
BCAT1 expression levels correlate with glioblastoma cell states in human tumors. (**A**) TCGA-GBM sample stratification based on BCAT1 expression into BCAT1^high^ (red), BCAT1^intermediate^ (gray) and BCAT1^low^ (blue). (**B**) Multidimensional scaling (MDS) analysis of BCAT1^high^ and BCAT1^low^ samples showing distinct clustering. (**C**) Pre-ranked gene set enrichment analysis (GSEA) showing an enrichment (Normalized enrichment score, NES) for neuronal-like glioblastoma signatures (blue) in BCAT1^low^ tumors (negative NES) and mesenchymal enrichment (red) in BCAT1^high^ (positive NES) samples. Only significant enrichments (*p* < 0.05) are shown. Signatures not associated with mesenchymal or neuronal subtypes are shown in gray. (**D**) Spearman’s correlation of BCAT1 expression (*x*-axis) and CIBERSORTx cell scores (*y*-axis) shows a positive correlation with the malignant cell score (*R* = 0.44, *p* = 8.1*10^-8^, left) and negative correlations with the macrophage (*R*=-0.33, *p* = 6.7*10^-5^, middle) and CD8-T-cell scores (*R* = -0.3, *p* = 0.00029, right).

Pre-ranked gene set enrichment analysis (GSEA) of differentially expressed genes (DEGs) of the BCAT1^high^ vs BCAT1^low^ groups using published glioblastoma molecular subtype signatures^31–34, 42^ showed significant enrichment of mesenchymal signatures in BCAT1^high^ tumors (Figure 1C, Positive normalized enrichment score (NES)). On the other hand, the BCAT1^low^ tumors showed strong enrichment in signatures associated with a neuronal-like phenotype, such as the proneural^31, 42^, neural progenitor cell (NPC) and developmental signatures^32–34^ (Figure 1C, Negative NES). Survival analysis of the complete TCGA-GBM cohort further showed that high BCAT1 expression in patients is associated with adverse prognosis (Supplementary Figure 1A). We further validated the cellular state findings in the patient glioblastoma stem cell line database HGCC^36^, where the mesenchymal and classical subtypes showed a higher BCTA1 expression than the proneural and neural-like (Supplementary Figure 1B). Using the highest and lowest BCAT1-expressing cell lines from this dataset based on z-score values, a single sample GSEA (ssGSEA) analysis showed clear clustering of the BCAT1-low cells according to proneural and developmental glioblastoma expressional programs whereas the ones with high BCAT1 expression showed mesenchymal or classical glioblastoma expression patterns (Supplementary Figure 1C and 1D).

Aggressive features of glioblastoma are associated with immunosuppression and an overall reduced CD8 T-cell infiltration^43^. As we observed that high BCAT1 expression correlates with a mesenchymal glioblastoma subtype, we used CIBERSORTx^44^ and a glioblastoma gene expression matrix^45^ to determine the cell-type composition of the TCGA samples. We found a highly significant positive correlation between BCAT1 expression and the malignant cell score, and a negative correlation for BCAT1 and the macrophage and CD8 T-cell scores (Figure 1D).

These results indicate a strong association between BCAT1 expression and the different molecular subtypes of glioblastoma, highlighting a more differentiated, neuronal-like expressional signature in BCAT1^low^ tumors. Furthermore, they imply a more prominent T-cell infiltration associated with low BCAT1 expression.

### Bcat1 knockout induces neuronal-like expression patterns in mouse and human glioblastoma cells

To address a potential causal link between BCAT1 expression and glioblastoma cell state, we used the mGB2 mouse glioblastoma neurosphere line derived from a *Trp53/Pten* double knockout genetic mouse model^24^ and the commonly used human glioblastoma line U251. Bcat1 was knocked out in mouse (Supplementary Figure 2A) and human glioblastoma cells, and a transcriptome analysis was performed.

In line with our observations in human samples, both the mGB2 Bcat1-KO and the U251 BCAT1-KO cells showed a strong enrichment of neuronal, whilst the respective Control cells showed mesenchymal state glioblastoma signatures (Figure 2A and Supplementary Figure 2B, respectively). Furthermore, gene ontology (GO) terms associated with neuronal processes were strongly enriched in both mGB2 and U251 BCAT1-KO cells (Supplementary Figure 2C and 2D, respectively). Notably, neurogenic differentiation 1 (*NeuroD1*), a master regulator of neurodevelopmental programs^46^, was among the top upregulated genes and a *NeuroD1*-target gene signature was highly enriched in mGB2 Bcat1-KO cells (Figure 2B).

**Figure 2.**
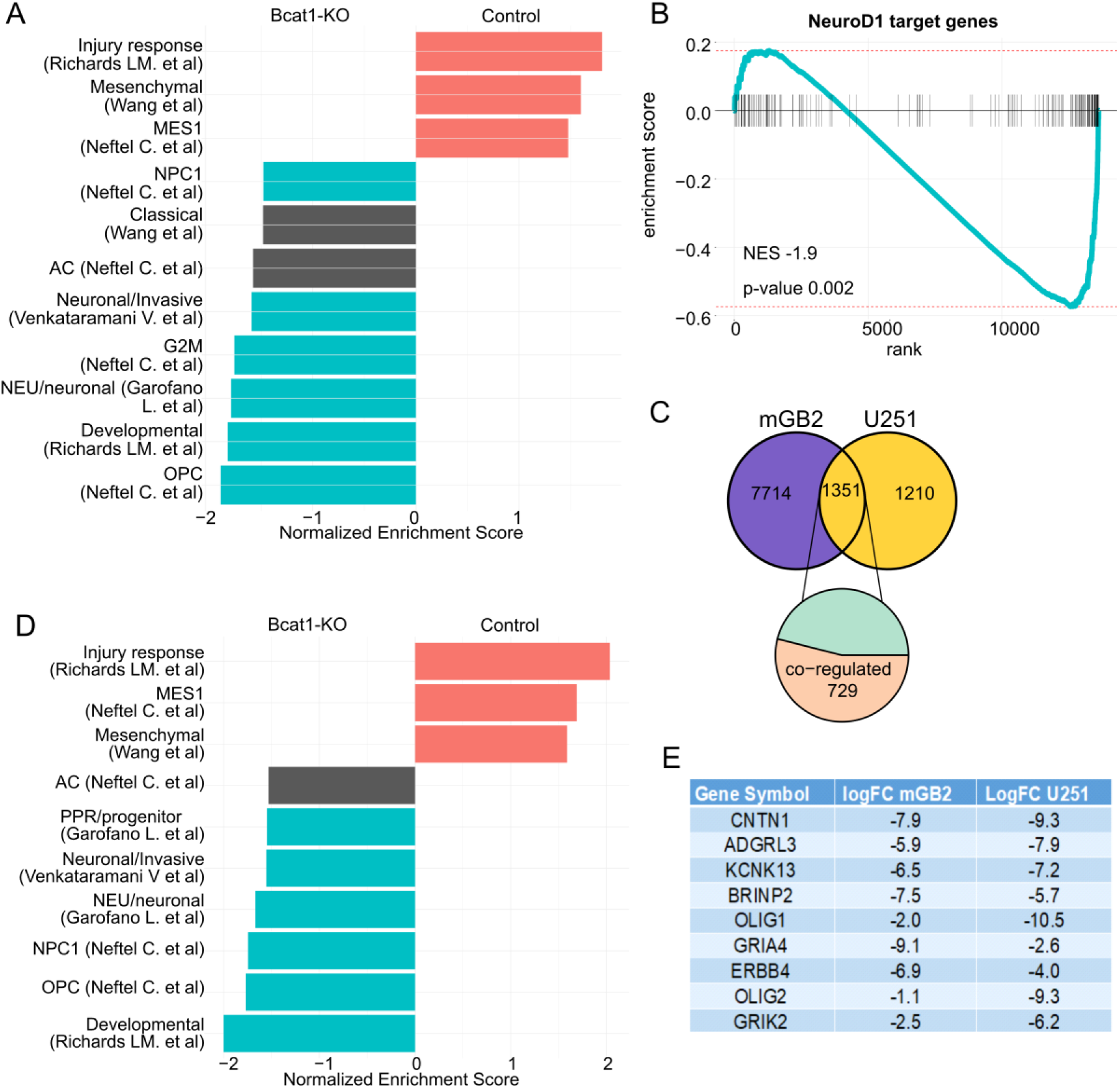
Bcat1 knockout induces a neuronal-like transcriptional shift. (**A**) Pre-ranked GSEA of DEGs between Control and Bcat1-KO mGB2 cells showing a strong enrichment of mesenchymal glioblastoma signatures (red) in Control cells (positive NES) and neuronal glioblastoma subtype signatures in Bcat1-KO cells (negative NES). Only statistically significant enrichments (*p* < 0.05) are shown. Signatures not associated with mesenchymal or neuronal subtypes are shown in gray. (**B**) Enrichment plot of genes regulated by NeuroD1 in the Bcat1-KO vs Control mGB2 cells (*p* = 0.002, *NES* = -1.9). (**C**) Venn-diagram of DEGs upon BCAT1 knockout and the overlapping genes between mGB2 (purple) and U251 (yellow). (**D**) Pre-ranked GSEA of the co-regulated genes in mGB2 and U251 cells upon knockout of BCAT1 showing a strong enrichment of mesenchymal signatures (red, positive NES) in Control and neuronal-like glioblastoma signatures (blue, negative NES) in BCAT1-KO cells. (**E**) Log fold change (logFC) of the top genes overexpressed in Bcat1-KO mGB2 and U251 cells involved in neuronal signaling and specification.

The similarity of transcriptional changes induced by BCAT1 knockout in the mouse and human lines, prompted us to identify genes commonly regulated by BCAT1. 729 genes were significantly co-regulated by BCAT1-KO (Figure 2C, “co-regulated”). A pre-ranked GSEA showed that these genes alone were sufficient to distinguish the Control and BCAT1-KO cells along the mesenchymal-neuronal differentiation axis (Figure 2D). Moreover, among the genes with the highest average upregulation in mGB2 and U251 BCAT1-KO cells, were mainly neuronal markers, or genes associated with neurodevelopmental processes (Figure 2E).

In summary, these data suggest that high BCAT1 expression maintains a poorly differentiated, mesenchymal cell state and that the lack of BCAT1 induces a transition to a neuronal-like expression pattern in human and mouse glioblastoma cells.

### Bcat1-KO mGB2 cells are prone to differentiation and differentiation-induced cell cycle arrest

mGB2 cells are regularly cultured as neurospheres in serum-free, stem-cell-supporting conditions. Therefore, we used this model to study the effects of Bcat1-KO on induced differentiation. To do this, we cultured the cells for 8 days either in stem cell (SC) conditions or in the presence of 5% fetal calf serum (FCS) as a non-specific differentiation-inducing agent. Treatment with FCS induced remarkable morphological differences between Control and Bcat1-KO mGB2 cells. Notably, control cells maintained a rounded shape with minimal cell extensions, while Bcat1-KO cells exhibited a highly elongated shape with pronounced cell extensions positive for tubulin beta 3 class III (Tubb3) (Figure 3A and Supplementary Figure 3A).

**Figure 3.**
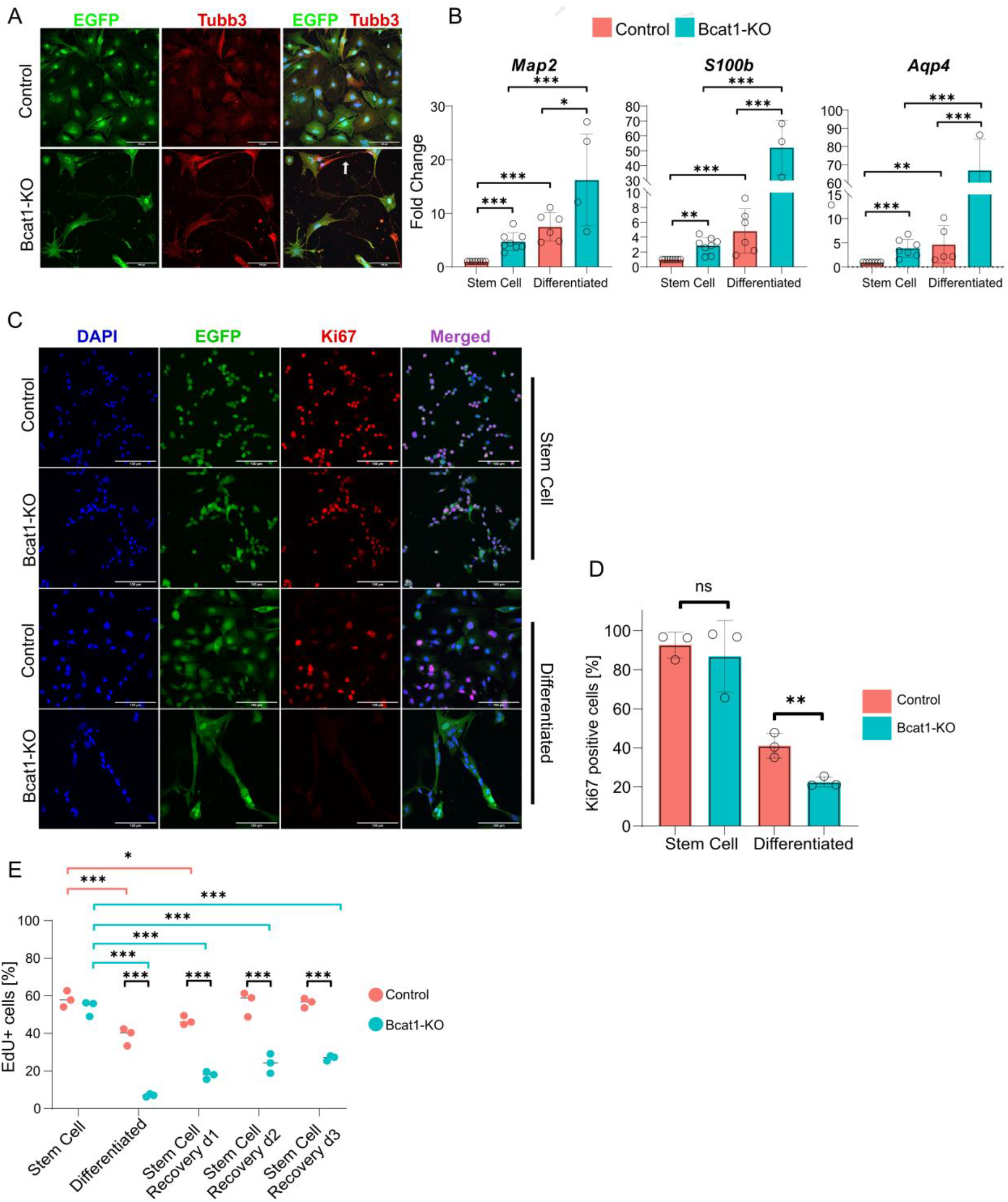
Bcat1-KO cells show a strong differentiation and are more prone to differentiation-induced cell cycle arrest *in vitro*. (**A**) Immunofluorescence of differentiated Control and Bcat1-KO mGB2 cells against EGFP (green), and Tubb3 (red). White arrow point to Tubb3-positive cellular extension indicative of neuronal differentiation. Scale bar = 100µm. (**B**) RT-qPCR quantification of differentiation genes in Control (red) and Bcat1-KO cells (blue) under Stem Cell or Differentiated conditions, showing increased expression in Bcat1-KO cells. Statistical comparisons were done using a one-way ANOVA with pairwise comparisons using Tukey’s multiple comparison test (*n* ⩾ 4). Error bars represent standard deviation. Only statistically significant comparisons are marked with stars. (**C**) Immunofluorescence against EGFP (green) and Ki67 (proliferation marker, red) in mGB2 cells under Stem Cell or Differentiation conditions, showing reduced Ki67 positivity in Bcat1-KO cells cultured under Differentiation conditions (**D**). Scale bar = 100µm. Statistical comparisons were performed using an unpaired, two-tailed student’s t-test. Error bars represent standard deviation (*n* = 3). (**E**) Click-iT EdU proliferation assay of Control (red) and Bcat1-KO (blue) cells cultured in Stem cell, Differentiation, or recovery (Stem Cell Recovery 1-3) conditions, showing a strong reduction of proliferation in Bcat1-KO cells in Differentiation conditions that is not fully recovered. Statistical analysis was performed using a two-way ANOVA test with Tukey’s multiple testing. Only statistically significant comparisons are marked with stars ns – non-significant, * – *p* ⩽ 0.05, ** – *p* ⩽ 0.01, *** – *p* ⩽ 0.001.

We then evaluated the expression of neuronal and glial differentiation markers by real-time quantitative PCR (RT-qPCR). As expected, FCS induced the expression of the glial and neuronal differentiation markers *Gfap* and *Tubb3*, respectively (Supplementary Figure 3B). In SC conditions, the expression of glial and neuronal markers *Aqp4*, *S100b* and *Map2* was already significantly higher in the Bcat1-KO cells (Figure 3B). In FCS conditions, the expression of these genes increased in both genotypes, but was strikingly higher in the differentiated Bcat1-KO than in Control cells (Figure 3B), indicating a stronger differentiation phenotype.

Next, we studied whether the pronounced differentiation in Bcat1-KO cells impacted cell proliferation. We cultured mGB2 Control and Bcat1-KO cells under SC or FCS conditions and quantified actively cycling cells using immunostaining against Ki67 (Figure 3C). Under SC conditions, most Control and Bcat1-KO cells were Ki67 positive (Ki67^+^). Upon the addition of FCS, the number of Ki67^+^ cells was reduced in both lines, however, it was significantly lower in Bcat1-KO cells compared to Control cells (on average 41% and 22%, respectively, Figure 3D).

Finally, we performed a Click-iT EdU proliferation assay using flow cytometry. Consistent with the Ki67 data, the number of EdU^+^ cells upon differentiation was significantly reduced in the Bcat1-KO cells compared to the Controls (from 57% to 40% and 56% to 8%, respectively) (Figure 3E). Furthermore, upon restoring SC conditions after differentiation, the Control cells completely regained their proliferative capacity after 2 days, whilst the Bcat1-KO cells showed only partial recovery (Figure 3E). These changes were also evident on the morphological level (Supplementary Figure 3C).

These data indicate that Bcat1-KO mGB2 cells are more responsive to differentiation signals than Control cells, suggesting that the loss of Bcat1 can overcome the *Trp53/Pten* loss-induced differentiation resilience of our double knockout model^24, 47^. Furthermore, the susceptibility of the Bcat1-KO cells to differentiate resulted in a strikingly increased differentiation-induced cell cycle arrest.

### Bcat1 maintains methylation-mediated suppression of neuronal-fate genes

Cell differentiation and phenotypic plasticity are critically regulated by epigenetic modifications such as DNA methylation at CpG dinucleotides^48^. We previously provided evidence that BCAT1 promotes DNA hypermethylation in AML cells by limiting cellular αKG, an essential co-factor of the DNA demethylases TET methylcytosine dioxygenases^19^. Thus, we hypothesized that changes in the CpG-methylation landscape could control the neuronal-like differentiation of Bcat1-KO mGB2 cells. To test this hypothesis, we first used a nuclear αKG sensor^41^ to confirm that BCAT1-KO increases αKG levels in the cell nucleus, where TET-dependent CpG demethylation occurs (Supplementary Figure 4A). Subsequent DNA-methylation analysis of mGB2 cells grown under SC conditions revealed an overall decrease of the proportion of hypermethylated sites in Bcat1-KO cells compared to Controls (Figure 4A), consistent with increased demethylase activity. Furthermore, differential CpG island methylation and gene expression inversely correlated between the Control and Bcat1- KO mGB2 cells (Figure 4B, *R* = -0.38, *p* < 0.01), indicating a methylation-based regulation of gene expression. Similarly, decreased methylation was observed in U251 BCAT1-KO compared to Control cells (Supplementary Figure 4B).

**Figure 4.**
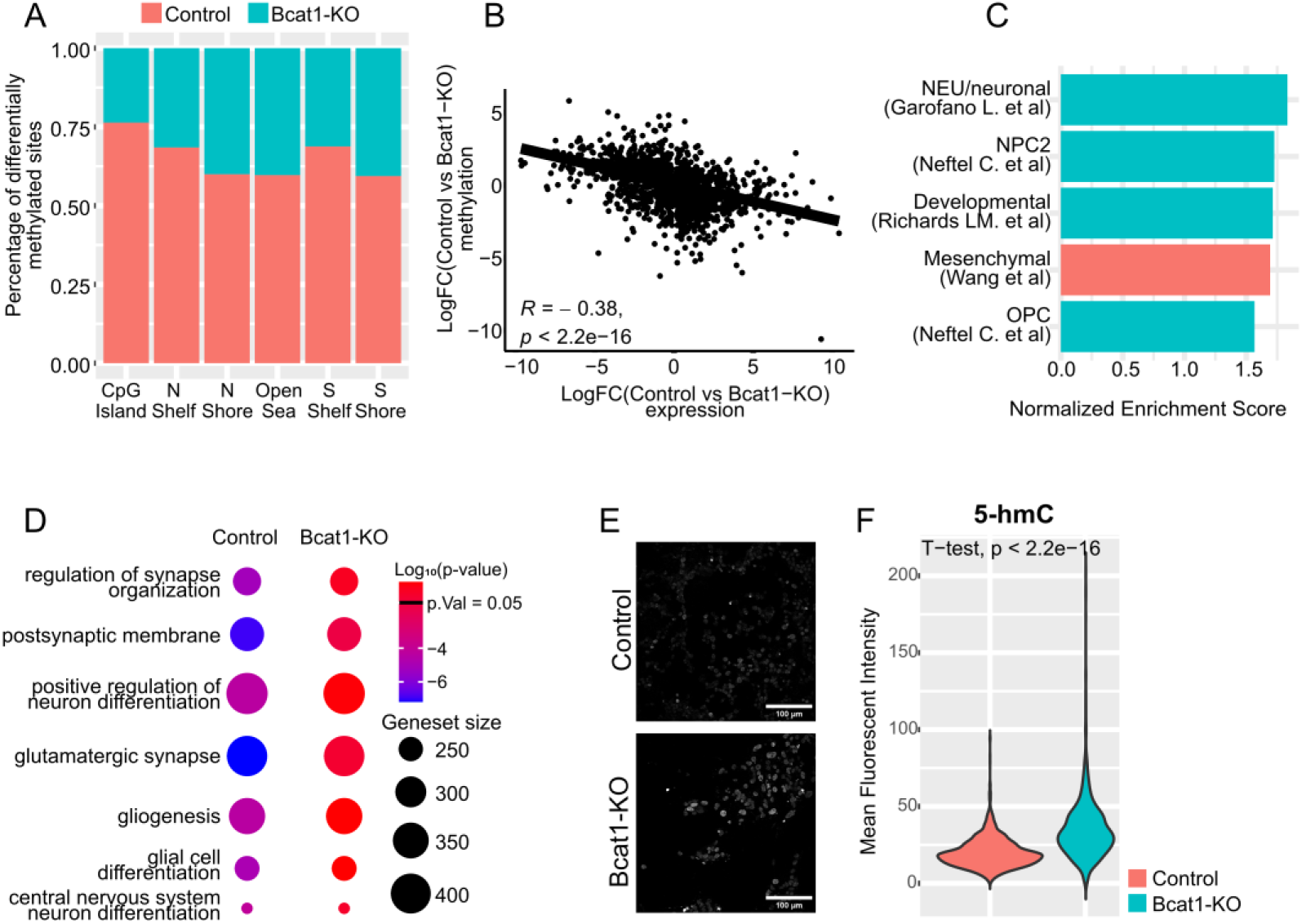
Bcat1 expression regulates DNA methylation. (**A**) Bcat1-KO mGB2 cells show an overall decrease in CpG methylation (blue) in comparison to the Control (red) including island-adjacent regions (S and N shelf and shores). (**B**) Spearman’s correlation of DEGs (*x*-axis) and differential gene methylation (*y*-axis) (*R* = -0.38, *p* < 2.2 * 10^-16^). (**C**) Robust rank aggregation analysis using hypermethylated CpG sites between Control and Bcat1-KO cells showing enrichment (positive NES) of genes regulating neuronal glioblastoma subtypes (blue) and a single mesenchymal signature (red) in Control cells. Only statistically significant (*p* < 0.05) enrichments are shown. (**D**) methylGSA enrichment of hypermethylated CpGs in Control and Bcat1-KO cells using neuronal differentiation GO-terms. Color scale represents the log_10_(*p*) of the enrichments and the size of each dot is proportional to the size of the geneset. (**E**) Immunofluorescent staining of 5-hmC in differentiated Control and Bcat1-KO mGB2 cells. Scale bar = 100µm. (**F**) Mean fluorescent intensity of the 5-hmC signal per nucleus of the differentiated Control (red) and Bcat1-KO (blue) mGB2 cells (*n* > 500 nuclei per condition). Statistical analysis was performed using an unpaired, two-tailed student’s t-test (*p* < 2.2 * 10^-16^). Experiment was repeated 3 times independently with the same outcome.

To test whether CpG methylation can account for repressing differentiation-related gene expression in the presence of Bcat1, we used differentially methylated CpG sites in a robust rank aggregation analysis^40^ against glioblastoma molecular subtype signatures. We found high enrichments for neuronal and developmental glioblastoma subtype gene sets in mGB2 Control cells (Figure 4C), but no significant enrichment in Bcat1-KO cells. Furthermore, Control cells showed a highly significant enrichment of non-cancer related neuronal differentiation signatures (Figure 4D) in a methylGSA analysis, suggesting that Bcat1 mediates the methylation-dependent suppression of neuronal fate genes.

To confirm that the altered methylation patterns observed upon Bcat1 knockout were a result of increased TET activity, we quantified the amount of 5-hydroxymethylcytosine (5-hmC), a direct product of TET-dependent 5-methylcytosine (5-mC) oxidation, in mGB2 Control and Bcat1-KO cells grown in FCS conditions (Figure 4E). Quantitative immunofluorescence analysis revealed significantly elevated levels of 5-hmC in the DNA of the Bcat1-KO cells, consistent with increased activity of TET enzymes (Figure 4F). Furthermore, the knockdown of either the Tet1 (shTet1) or Tet3 (shTet3) genes in the Bcat1-KO mGB2 cells (Supplementary Figure 5A and 5B) resulted in reduced expression of neuronal differentiation genes such as *NeuroD1* (Supplementary Figure 5C). A more detailed analysis by methylation array showed shTet1 and shTet3 efficiently restored the enrichment of hypermethylation of neuronal and glial differentiation-associated genes in comparison to the control (shLuc) Bcat1-KO cells (Supplementary Figure 5D).

In summary, these analyses reveal that Bcat1 knockout increases TET activity, resulting in an overall decrease in DNA methylation that strongly correlates with the expression of differentiation-related genes. These findings suggest a mechanistic explanation for BCAT1’s function in glioblastoma cell plasticity, wherein BCAT1 restricts cellular αKG resulting in a hypermethylation and repression of neuronal fate genes.

### Bcat1-KO induces glioblastoma cell differentiation and promotes an anti-tumor immune response *in vivo*

To test whether Bcat1-KO cells are susceptible to differentiation *in vivo*, we orthotopically transplanted Bcat1-KO and Control mGB2 cells in immunocompetent C57BL/6 mice and characterized the tumor phenotypes at early (4 weeks) and advanced (upon reaching termination criteria) time points.

EGFP immunostaining detected signs of cell differentiation in Bcat1-KO cells already at week 4 (Figure 5A). These differentiation features became more pronounced at the advanced stage with all Bcat1-KO cells showing one or multiple prolonged cell extensions reminiscent of neuronal protrusions (Figure 5B). In contrast, Control mGB2 cells exhibited the typical glioblastoma morphology, appearing as tightly packed rounded tumor cells at both early and late time points (Figure 5A and 5B). These observations were supported by a reduced expression of the glioblastoma stem cell marker Sox2 in the Bcat1-KO mGB2 tumors, further confirming cellular differentiation in these cells (Supplementary Figure 6A and 6B).

**Figure 5.**
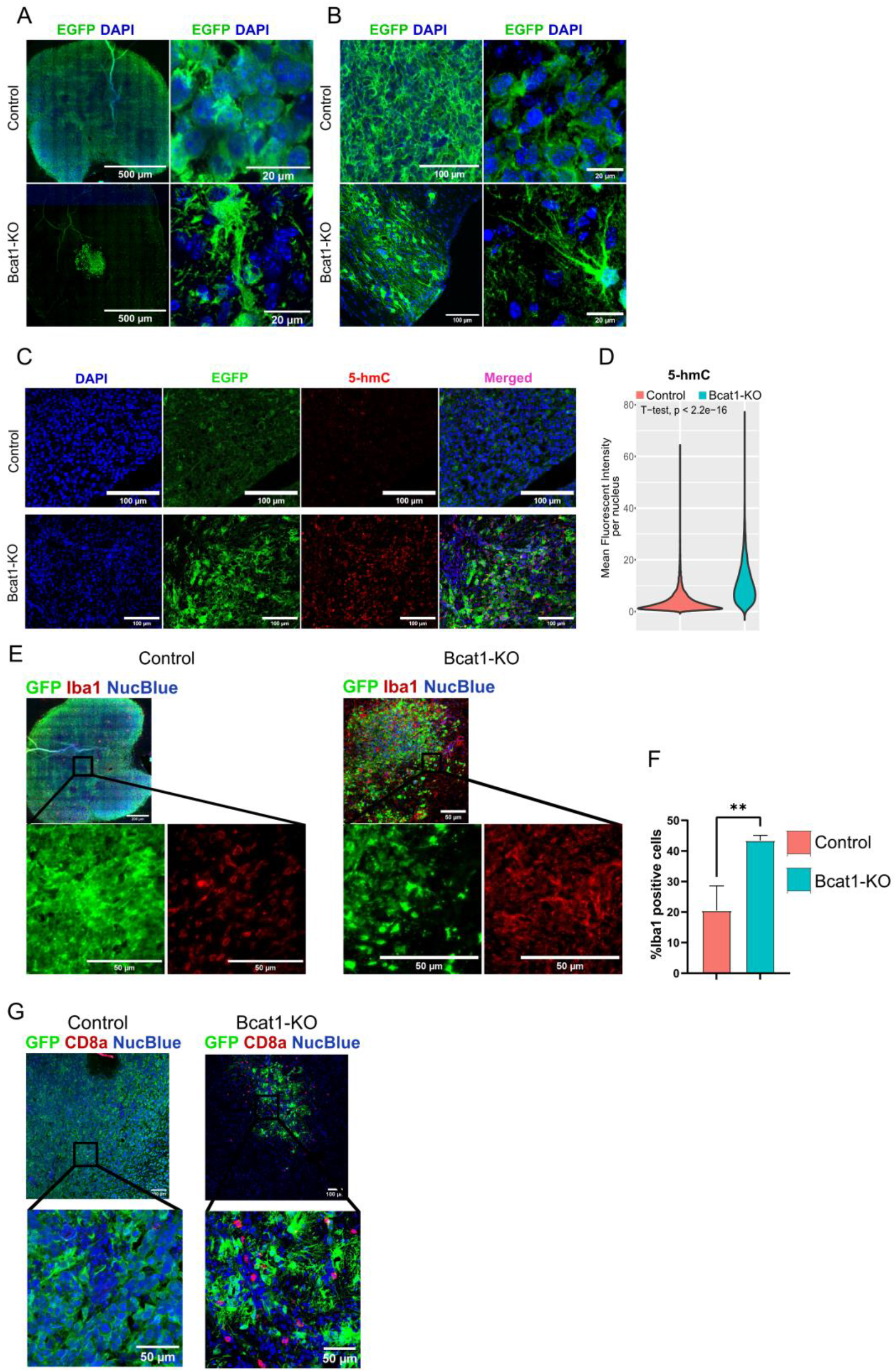
Bcat1 knockout induces tumor cell differentiation in an immunocompetent mouse model. (**A**) Tumor immunofluorescent imaging (left) and cellular morphology confocal microscopy (right) of 4-week and (**B**) advanced Control and Bcat1-KO tumors with anti-EGFP labeling (green). Scale bar = (**A**) 500µm (left) and 20µm (right) and (**B**) 100µm and 20µm. (**C**) Immunofluorescent labeling of 5-hmC (red) in Control and Bcat1-KO tumors 4 weeks post-injection labeled against EGFP (green, tumor cells). DNA was labeled using DAPI (blue). Scale bar = 100µm. (**D**) Mean fluorescent intensity of the 5-hmC signal per nucleus in Control (red, *n* = 3) and Bcat1-KO (blue, *n* = 3) tumors (more than 200 nuclei were quantified per mouse). Statistical analysis was performed using an unpaired, two-tailed student’s t-test (*p* < 2.2 * 10^-16^). (**E**) Immunofluorescent saining against EGFP (green) and Iba1 (myeloid marker, red) of Control (left) and Bcat1-KO (right) tumors 4 weeks post-injection with cut-outs highlighting signs of differentiation and increased myeloid density in Bcat1-KO tumors. Scale bar = 200µm and 50µm (Control), and 50µm (Bcat1-KO). (**F**) Quantification of Iba1^+^ cells in the tumor region of Control (red, *n* = 3) and Bcat1-KO (blue, *n* = 3) tumors. Error bars represent standard deviation. Statistical comparison was performed using a two-tailed unpaired student’s t-test. (**G**) Immunofluorescent staining against EGFP (green) and CD8a (red) in advanced Control and Bcat1-KO tumors with representative cut-outs. Scale bar = 100µm and 50µm. NucBlue or DAPI were used to stain DNA (blue) in all images. ns – non-significant, * – *p* ⩽ 0.05, ** – *p* ⩽ 0.01, *** – *p* ⩽ 0.001.

In order to examine whether this differentiation phenotype is accompanied by an increase in 5-hmC, we used quantitative immunofluorescence imaging of the early-stage Control and Bcat1-KO tumors (Figure 5C). The analysis showed a significant (*p <* 0.01) increase in 5- hmC in Bcat1-KO tumor cells (Figure 5D), indicating that the lack of Bcat1 results in a higher TET activity *in vivo*, consistent with our *in vitro* findings (Figure 4F).

As the tumor immune microenvironment plays a crucial role in regulating tumor growth, and our analysis of patient tumors revealed that higher BCAT1 expression correlates with lower effector T-cell infiltration (CD8 T-cells), we proceeded to analyze the immune composition of the mouse tumors.

Early in tumor development, we observed the expected myeloid morphology of glioblastoma-associated myeloid cells in Control tumors (Figure 5E, Control). Remarkably, in the Bcat1- KO tumors we observed a significant increase in the myeloid population accompanied by altered morphological features (Figure 5E, Bcat1-KO), which was even more pronounced at advanced time point (Supplementary Figure 6C). Quantification of Iba1-positive cells identified more than double the average number of myeloid cells in the Bcat1-KO tumors (Figure 5F). Next, we visualized the CD8-positive cell population in the tumor sections by immunofluorescent staining. At an early time point, tumors of both genotypes showed signs of T-cell infiltration with a tendency towards increased infiltration in the Bcat1-KO mice (Supplementary Figure 6D). However, at the advanced stage of tumor development very few CD8 T-cells were detected in the Control tumors, whereas the infiltration remained high in Bcat1-KO tumor residuals (Figure 5G, Supplementary Figure 6E).

These data show that Bcat1 knockout-induced glioblastoma differentiation occurs early during tumor development *in vivo*. Furthermore, Bcat1-KO appears to promote the remodeling of the tumor immune microenvironment resulting in a persistent infiltration of CD8 T-cells, consistent with our observations in the BCAT1^low^ human glioblastoma samples.

### Bcat1-KO significantly prolongs survival time but does not completely abrogate tumor growth in immunodeficient mice

To dissect the impact of the two Bcat1-dependent phenotypes, immunosuppression and tumor cell differentiation, on mouse survival, we transplanted Control and Bcat1-KO mGB2 cells into immunodeficient NOD SCID gamma (NSG) mice, lacking both the myeloid and lymphocytic immune compartments. Unlike in immunocompetent mice^25^, Bcat1-KO cells grew detectable tumors in NSG mice (Figure 6A). However, animal survival was still significantly longer than for mice transplanted with Control cells (Figure 6B, *p* = 0.0007). Compared to a complete lack of tumor growth of Bcat1-KO cells in immunocompetent mice (Figure 6B, dashed lines), these results highlight the importance of both so-far-described phenotypes: the immune system is necessary to completely abrogate tumor growth of the Bcat1-KO cells and the immune-independent differentiation effects significantly contribute to slower tumor growth and prolonged survival. To distinguish weather the myeloid or lymphocytic compartments were crucial for Bcat1-KO tumor suppression, we injected tumor cells into T cell deficient Rag2-KO mice which lack functional T cells but maintain myeloid cells such as microglia and macrophages. Here we observed the same tumor growth pattern as in the NSG mice, strongly suggesting that T cells are necessary for the suppression of Bcat1-KO tumor growth (Supplementary Figure 6F).

**Figure 6.**
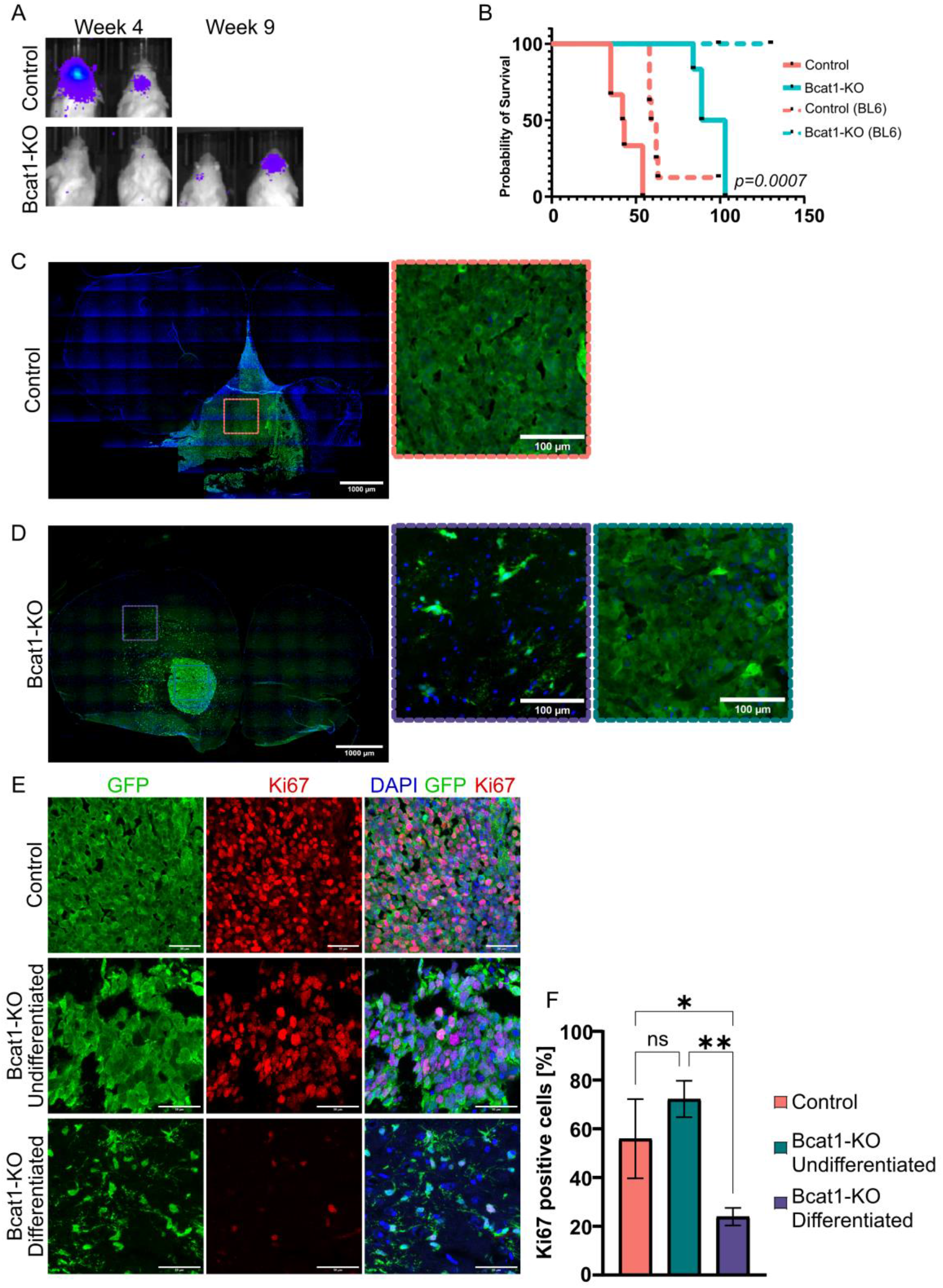
Bcat1-KO cells form tumors with highly differentiated regions and obvious cell cycle arrest in immunocompromised mice. (**A**) IVIS of Luciferase-expressing Control and Bcat1-KO tumors 4 weeks and 9 weeks post-injection. (**B**) Kaplan-Meier survival analysis of NSG mice injected with Control (full red line, *n* = 6) and Bcat1-KO (full blue line, *n* = 6) mGB2 cells. Previously published^25^ survival times of C57BL/6 mice transplanted with Control (dashed red line, *n* = 8) and Bcat1-KO (dashed blue line, *n* = 8) cells are shown for comparison. (**C**) Whole section immunofluorescent imaging of Control and (**D**) Bcat1-KO tumors in the NSG mice labeled with EGFP (green). Control tumors show poorly differentiated morphology (red box), while Bcat1-KO tumors present with highly differentiated (purple box) and poorly differentiated (blue box) regions. Scale bar = 1000µm and 100µm. (**E**) Confocal immunofluorescence images of Control tumors and differentiated and undifferentiated regions of Bcat1-KO tumors labeled with anti-EGFP (green) and anti- Ki67 (red). Scale bar = 50µm. (**F**) Quantification of Ki67+ cells in the Control tumors (red, *n* = 3) and undifferentiated (blue, *n* = 3) or differentiated (purple) regions of Bcat1-KO tumors. More than 100 cells were quantified per condition per mouse. Statistical comparisons were performed using a one-way ANOVA analysis with Tukey’s multiple testing. Error bars represent standard deviation. ns – non-significant, * – *p* ⩽ 0.05, ** – *p* ⩽ 0.01, *** – *p* ⩽ 0.001.

Using anti-EGFP fluorescent labeling in sections form mGB2 tumors grown in NSG mice, we observed the typical poorly differentiated phenotype of the Control cells (Figure 6C). In contrast, the highly differentiated morphological features of the Bcat1-KO cells (Figure 6D, purple insets) matched those found in the immunocompetent model (Figure 5C). Interestingly, after examining the whole coronal sections of the Bcat1-KO tumors in immunodeficient mice, we also observed small undifferentiated regions (Figure 6D, blue inserts), which were not apparent in C57BL/6 mice (Figure 5D).

To test whether the morphologically distinct population of Bcat1-KO tumor cells also showed different proliferative capacities, we quantified the portion of Ki67^+^ cells in differentiated and undifferentiated regions of the Bcat1-KO and in Control tumors (Figure 6E). We observed a significant reduction of the number of Ki67^+^ cells in the differentiated regions of the Bcat1- KO tumors compared to both Control and Bcat1-KO undifferentiated regions (Figure 6F), suggesting that the remaining small undifferentiated regions primarily drive Bcat1-KO tumor growth.

In summary, these data indicate that the immune system plays a role in abolishing Bcat1-KO tumor growth, evidenced by the growth Bcat1-KO tumors in immunodeficient NSG and Rag2-KO mice. However, Bcat1-knockout-induced differentiation and cell cycle arrest occur independent of the host’s immune system and result in the doubling of the survival time. These results highlight the importance of Bcat1 in preventing tumor cell differentiation and promoting glioblastoma aggressiveness through maintaining immunosuppression.

## Discussion

Glioblastoma is characterized by its extensive intra-tumoral heterogeneity, coupled with pronounced immunosuppression and phenotypic plasticity, allowing individual tumor cells to transition between at least four major cell states to evade therapeutic pressure^31, 33, 34^. Glioblastoma cells have been shown to differentiate to various extents both *in vitro* and *in vivo*^8, 12, 49^. A striking example was recently provided by studies showing that partial neuronal-like differentiation is co-opted by glioblastoma cells to promote migration through the brain utilizing neuronal signaling pathways. This process is used by glioblastoma cells in order to revert back to their proliferative and poorly differentiated state once the differentiation cues subside^8, 12^.

We previously reported a significant reduction of glioblastoma growth upon BCAT1 knockdown using xenograft mouse models^17, 25^. Here, we provided evidence indicating BCAT1 as a regulator of glioblastoma cell differentiation and phenotypic plasticity. Using human patient data and a glioblastoma stem cell database, we show that low BCAT1 expression correlates with a more pronounced neuronal and differentiated features of tumors. Furthermore, knocking out Bcat1 in a primary mouse glioblastoma stem cell model that faithfully recapitulates human glioblastoma^24^, or in a human glioblastoma cell line, induced an even more striking shift towards a differentiated state. *In* vivo, using a syngeneic mouse model, we show that Bcat1-KO cells not only differentiate but also fail to develop into tumors. In immunodeficient mice, we demonstrate that even though Bcat1-KO-induced differentiation remains a major contributing factor to prolonged survival, the presence of a functional immune system, and in particular the lymphoid compartment, is necessary to completely abrogate the growth of these cells.

Inducing the differentiation of glioblastoma cells into either neuronal or glial lineages has been a target of investigation for the potential treatment of glioblastoma and other brain tumor entities^50, 51^. In our study, we show that human and mouse Bcat1-KO cells overexpress major neuronal fate regulators such as *NeuroD1* and *Ascl1*, both of which have been shown to cause terminal differentiation and neuronal fate specification in glioblastoma cell lines and mouse models^52, 53^. These findings indicate that similar differentiation effects can be achieved without the need for ectopic overexpression, through DNA hypomethylation of neuronal differentiation genes associated with the downregulation of BCAT1.

Relative depletion of αKG has been shown to play a central role in maintaining the undifferentiated state of pluripotent stem cells^54^. Furthermore, suppression of αKG-dependent processes through competitive binding of the oncometabolite 2-hydroxyglutarate has been implicated in the pathomechanism of IDH-mutant gliomas^55, 56^. Related mechanisms have been proposed to explain the oncogenic nature of mutations in the Krebs cycle enzymes succinate dehydrogenase and fumarate hydratase^57, 58^. In this study, we propose αKG depletion by BCAT1 and consequential hypermethylation and transcriptional repression due to reduced TET-demethylase activity as a mechanism of maintaining a mesenchymal-like cell state in glioblastoma. We previously demonstrated the prognostically relevant association of BCAT1 expression and DNA methylation in AML^19^ and thus propose that the outlined BCAT1-dependent mechanisms might also regulate cell state transitions in other cancers in which BCAT1 has been identified as a driver of tumor cell proliferation^17, 18^.

In summary, we propose a novel mode of action of BCAT1 in promoting glioblastoma tumorigenesis through αKG depletion and subsequent reduced activity of TET enzymes resulting in the suppression of differentiation-related genes. Furthermore, we show that BCAT1 expression contributes to the immunosuppressive nature of the tumor microenvironment. These findings implicate BCAT1 activity as a potential novel target for differentiation and immunomodulatory therapies in glioblastoma and other BCAT1- overexpressing tumor entities.

## Acknowledgements

The authors thank Petra Schroeter for her excellent technical assistance. We would also like to acknowledge the DKFZ Microscopy and Proteomics and Omics IT and Data Management core facilities for technical support.

## Funding

No funding was received towards this work.

## Competing interests

None declared.

## Notes

### Competing Interest Statement

The authors have declared no competing interest.

