## Supplementary Figures and Tables for "BCAT1 regulates glioblastoma cell plasticity and contributes to immunosuppression"

### Supplementary Figure 1

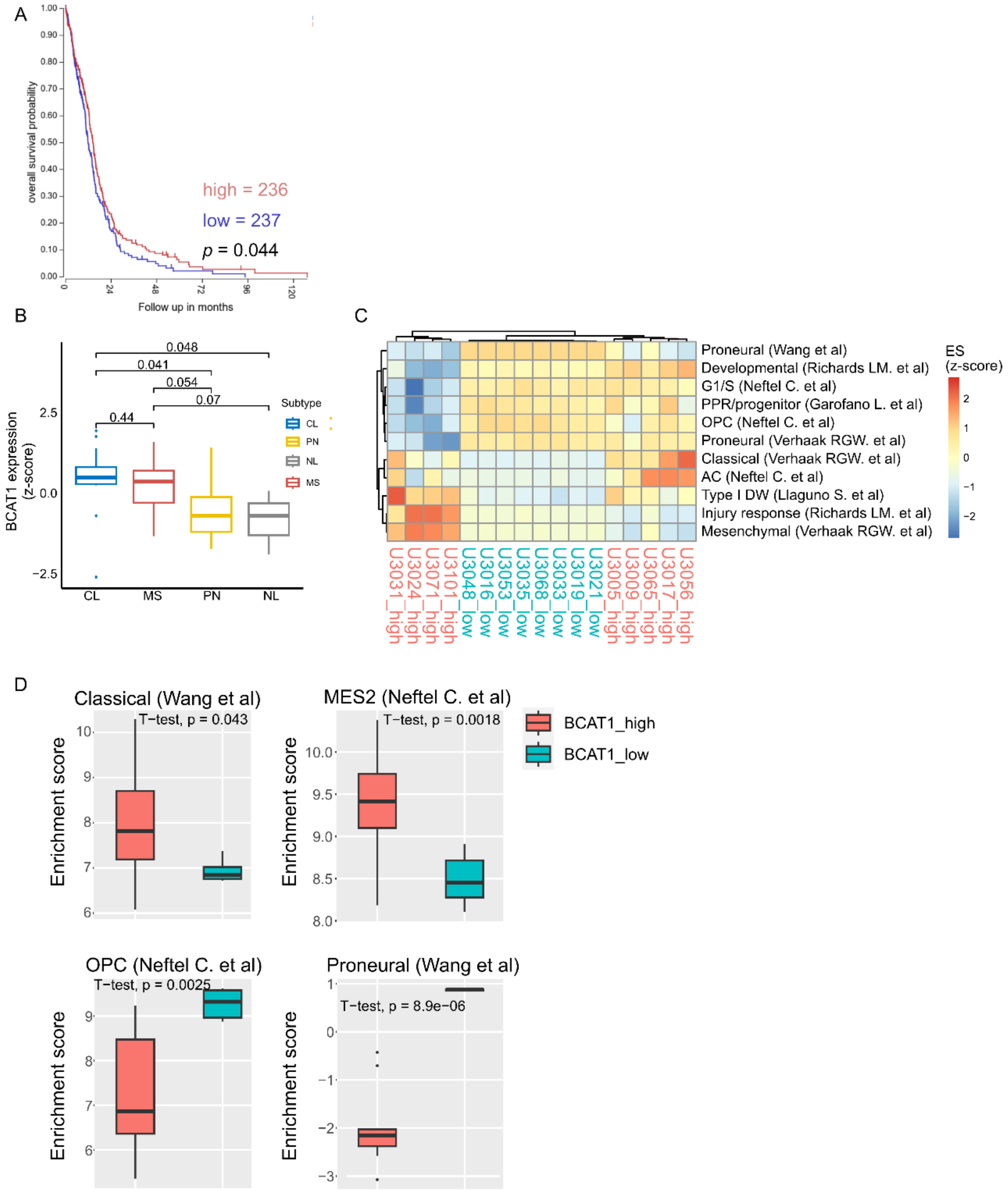


**Supplementary Figure 1 Low BCAT1-expression is associated with neural and proneural glioblastoma subtypes in patient tumors and patient-derived glioblastoma stem cell lines.** (A) (A) Kaplan-Meier survival curve of glioblastoma patients in the TCGA-GBM cohort stratified based on BCAT1 expression as high (red) and low (blue), relative to median expression. The curve was generated using the R2 platform. (B) HGCC dataset BCAT1 expression represented as z-scores according to glioblastoma line subtype (CL – classical, PN – proneural, NL – neuronal-like, MS – mesenchymal). Pair-wise statistical comparison was performed using unpaired, two-tailed student's t-test with significance values denoted above each comparison. (C) Enrichment score (ES) heatmap of single sample gene set enrichment analysis (ssGSEA) of BCAT1-high (red) and BCAT1-low (blue) HGCC cell lines (z-score > 1and z-score < -1, respectively) against published glioblastoma subtype signatures. ES are represented as z-score values. (D) Representative signature ESs averaged across BCAT1-high (red) and BCAT1-low (blue) HGCC cell lines. Statistical comparisons were performed using an unpaired, two-tailed student's t-test with significance levels denoted in each panel. Error bars signify 1.5 IQR.

### Supplementary Figure 2

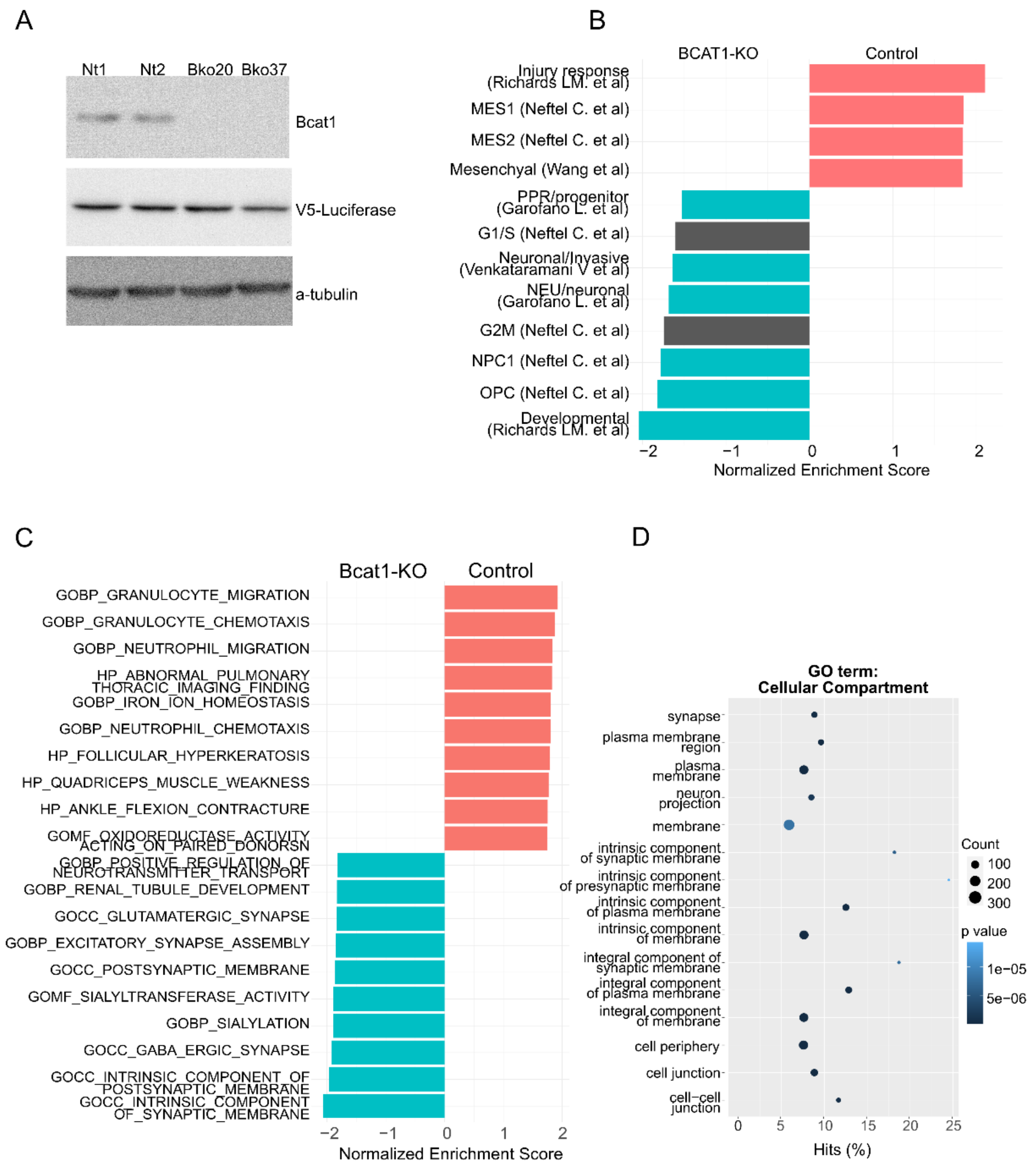


**Supplementary Figure 2 Characterization of Bcat1-KO mGB2 and U251 glioblastoma cells.** (A) Western blot confirmation of CRISPR/Cas9-mediated Bcat1-KO in mGB2 clones (Bko20 and Bko27) and confirmation of equal V5-Luciferase expression in Control clones (Nt1 and Nt2) and the 2 Bcat1-KO clones. αTubulin was used as a loading control. (B) Pre-ranked GSEA of DEGs between Control (positive NES) and BCAT1-KO (negative NES) U251cells against the glioblastoma molecular subtype signature set. Mesenchymal signatures are highlighted in red, neuronal signatures in blue and the ones not associated with either in gray. Only statistically significant signatures ($p < 0.05$) are represented. (C) Pre-ranked GSEA of DEGs between control (red) and Bcat1-KO (blue) mGB2cells against the BP ontology dataset. Only significant enrichments ($p < 0.05$) are shown. (D) GO-term enrichment analysis of genes overexpressed in BCAT1-KO U251 cells using Cellular Component ontology terms.

### Supplementary Figure 3

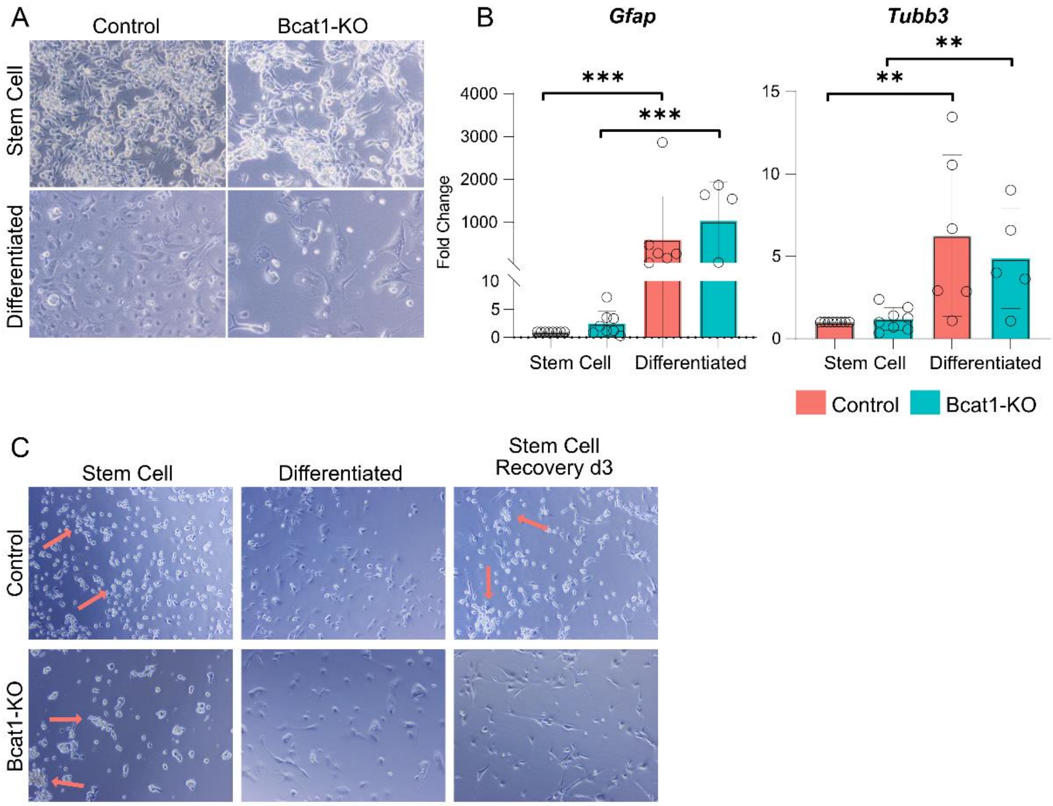


**Supplementary Figure 3 In vitro differentiation of mGB2 cells.** (A) Phase contrast microscopy demonstrating morphological features of Control and Bcat1-KO cells in Stem Cell and Differentiation (5% FCS) conditions. (B) RT-qPCR gene expression analysis of *Gfap* and *Tubb3* expression as markers of differentiation in Control (red) and Bcat1-KO (blue) cells in Stem Cell and Differentiation conditions. *Tbp* was used for obtaining the dCt values and all samples were normalized to the Control Stem Cell condition. Statistical analysis was performed using one-way ANOVA testing (n = 4-8) and Tukey's multiple comparison tests. Only significant comparisons ($p < 0.05$) are marked with stars. Error bars represent standard deviation. (C) Phase contrast microscopy demonstrating morphological features of Control and Bcat1-KO cells in Stem Cell and Differentiation conditions and upon 3 days of recovery after differentiation. Red arrows point towards a typical stem cell clustered phenotype. ns – non-significant, * – $p \leqslant 0.05$, ** – $p \leqslant 0.01$, *** – $p \leqslant 0.001$.

Supplementary Figure 4

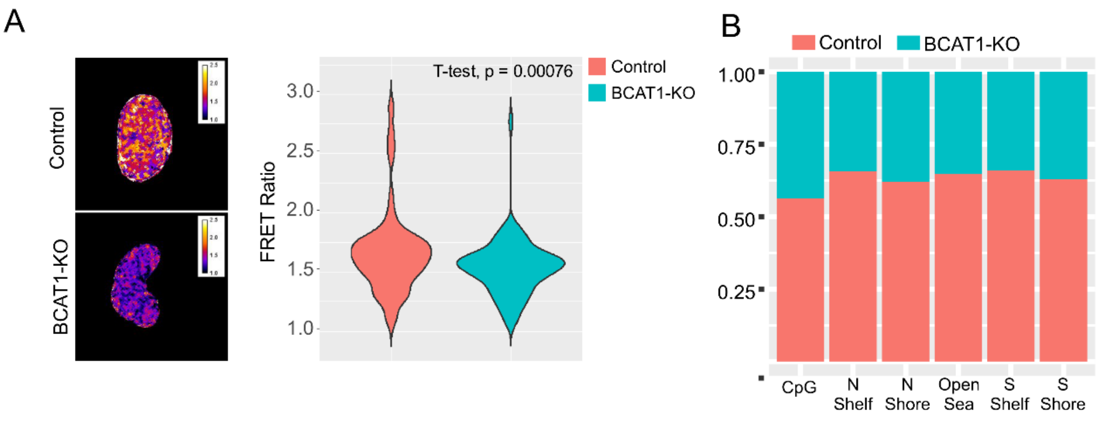


**Supplementary Figure 4 BCAT1-KO alters methylation patterns in glioblastoma cells.** (A) Representative radiometric images and quantification of the FRET ratio detected in the nuclei of Control (red, n = 140 nuclei) and BCAT1-KO (blue, n = 138 nuclei) U251 cells expressing the αKG-sensor. Statistical comparison was performed using an unpaired, two-tailed student's t-test. (B) Percentage of hypermethylated sites in Control (red) or BCAT1-KO (blue) cells according to probe localization in U251 cells. Sites with a difference in β-values higher than 20% between the two lines were used (n = 1 technical replicate). (C) Spearman correlation analysis of differential gene expression of neuronal genes form Figure 4D (x-axis) and differential gene methylation (y-axis) (R = -0.45, p = 1.2*10-8).

### Supplementary Figure 5

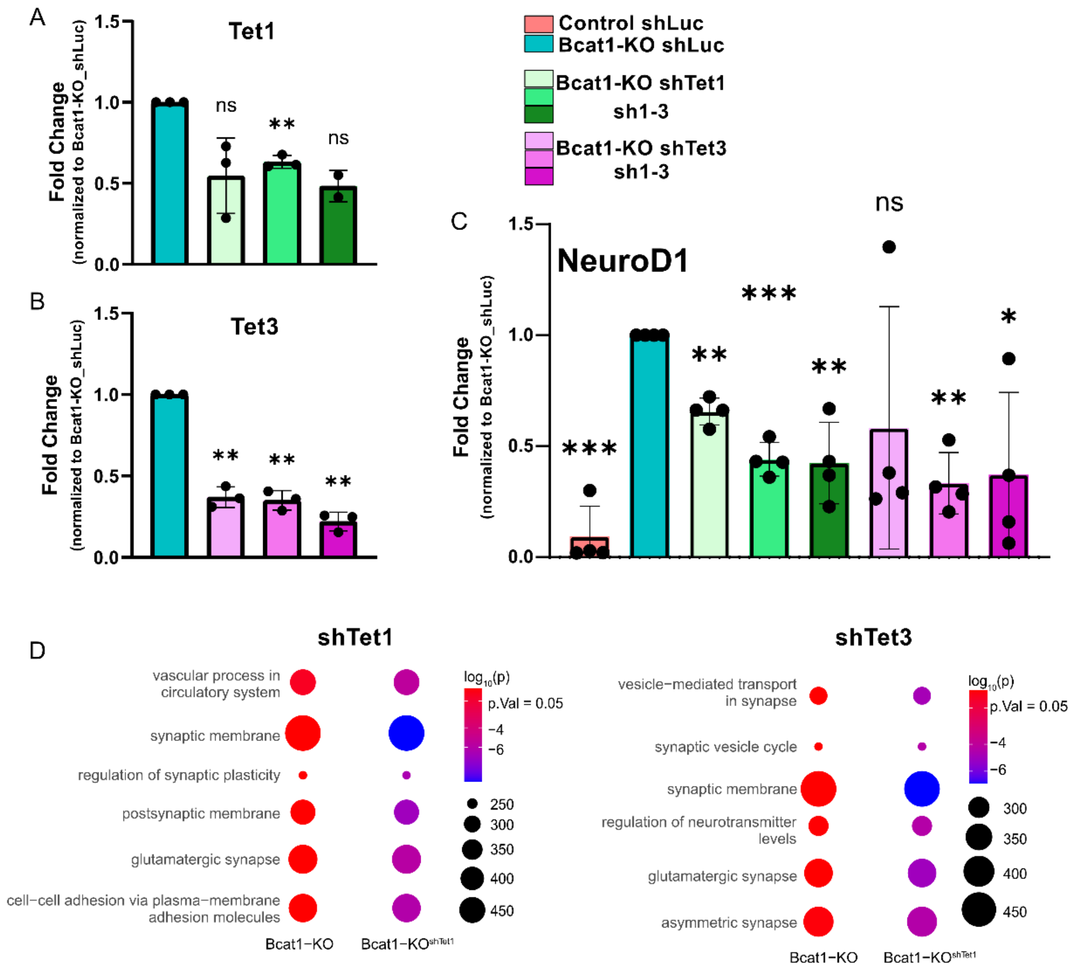


**Supplementary Figure 5 knockdown (KD) of Tet1 and Tet3 enzymes alters the expression of neuronal differentiation genes in mGB2 mouse glioblastoma cells**. Knockdown efficiency of Tet1 (A) and Tet3 (B) enzymes in Bcat1-KO mGB2 cells with 3 different shRNA constructs were confirmed with RT-qPCR expression analysis. shRNA against luciferase (shLuc) used as control. Statistical comparisons were performed using a single-column t-test relative to the control (Bcat1-KO shLuc). (C) Expression of NeuroD1 in Control and Bcat1-KO mGB2 cells as well as shTet1 1-3 (green) and shTet3 1-3 (purple) KD Bcat1-KO mGB2 cells. Statistical comparisons were performed using a single-column t-test relative to the control (Bcat1-KO shLuc). (D) Top enriched

GO-signatures using methylGSA enrichment analysis of hypermethylated CpGs in Tet1-KD (left) and Tet3-KD (right) mGB2 Bcat1-KO cells in comparison to the shLuc mGB2 Bcat1-KO controls. Color scale represents the $\log_{10}(p)$ of the enrichments and the size of each dot is proportional to the size of the gene set. 3 different shTet1 and shTet3 constructs were used for the methylation array together with 3 replicates of the Bcat1-KO mGB2 cells.

### Supplementary Figure 6

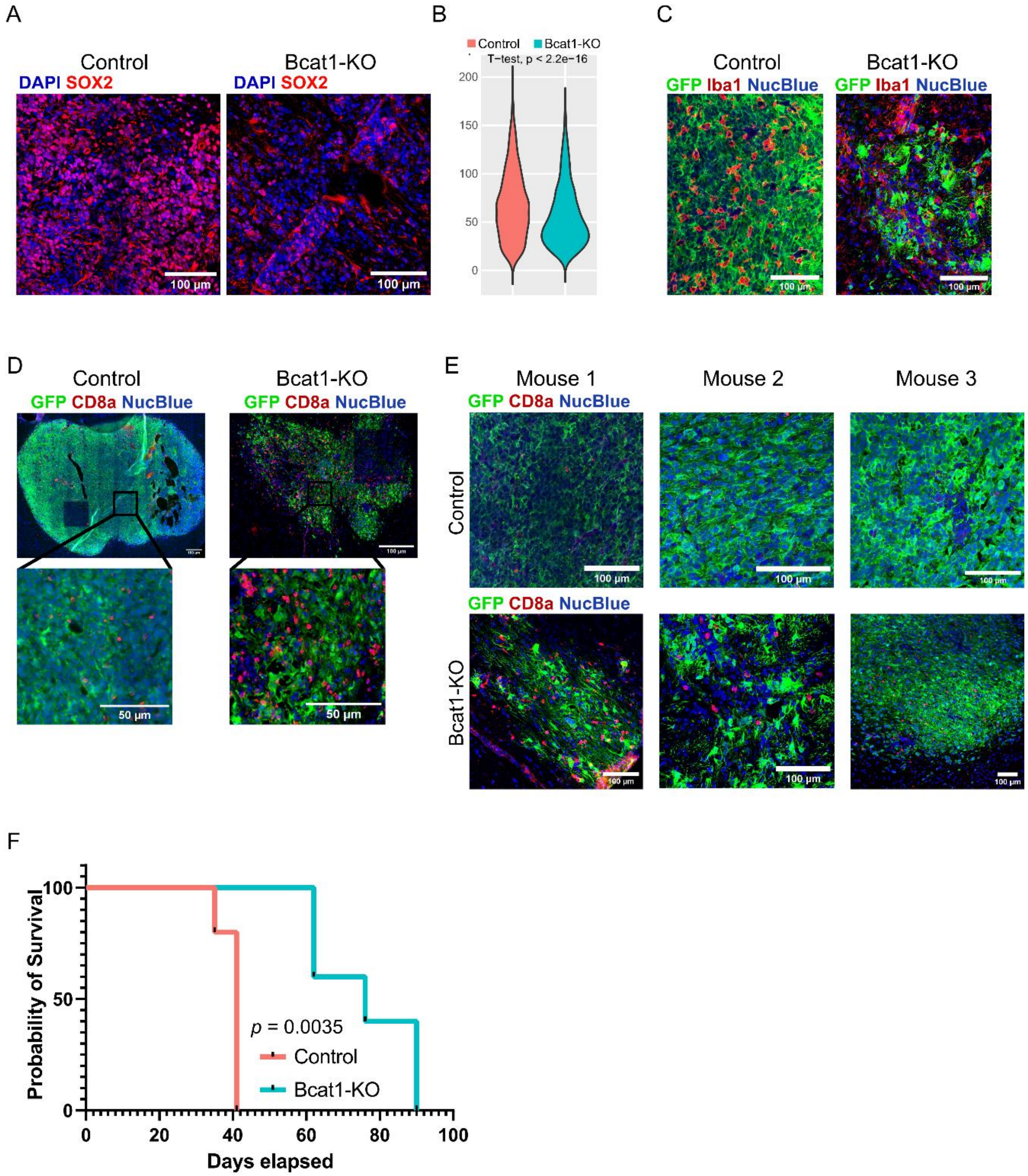


**Supplementary Figure 6 Bcat1 expression influences tumor cell differentiation and the immune microenvironment *in vivo*.** (A) Immunofluorescent labeling and confocal microscopy of Control and Bcat1-KO tumor cells against Sox2 (red). DNA was stained using DAPI. Scale bar

= 100μm. (B) Mean fluorescent intensity quantification of Sox2 expression per nucleus in Control and Bcat1-KO mice (n=3). Statistical analysis was performed using unpaired, two-tailed student's t-test and statistical significance is denoted on top of the panel. (C) Representative images of advanced tumor immunofluorescent labeling using EGFP (green) and Iba1 (red) in a Control and Bcat1-KO tumor. (D) Representative whole-tumor immunofluorescence of early Control and Bcat1-KO tumors, with a tendency towards increased CD8 T-cell infiltration (red) in the controls. EGFP was used to label tumor cells (green). Scale bar = 100μm in whole tumor imaging and 50μm in representative inserts. (E) Images of CD8 T-cell infiltration using immunofluorescence (CD8a, red) in 3 advanced Control and Bcat1-KO tumors. EGFP was used to label tumor cells (green). Scale bar = 100μm. DAPI was used for DNA staining (blue). (F) Kaplan-Meier survival curve of Rag2-KO mice injected with Control (red, n = 5) and Bcat1-KO (blue, n = 5) mGB cells. Significance was determined using log-rank analysis.

**Supplementary Table 1 List of primers used for the qPCR analysis**

| Oligo | Sequence |
|---|---|
| mAqp4 forward | ATCAGCATCGCTAAGTCCGTC |
| mAqp4 reverse | GAGGTGTGACCAGGTAGAGGA |
| mCspg4 forward | GCTGTCTGTTGACGGAGTGTT |
| mCspg4 reverse | CGGCTGATTCCCTTCAGGTAAG |
| mGfap forward | CGGAGACGCATCACCTCTG |
| mGfap reverse | CGGAGACGCATCACCTCTG |
| mMap2 forward | GCCAGCCTCGGAACAAACA |
| mMap2 reverse | GCTCAGCGAATGAGGAAGGA |
| mS100b forward | TGGTTGCCCTCATTGATGTCT |
| mS100b reverse | CCCATCCCCATCTTCGTCC |
| mTbp forward | ATGATGCCTTACGGCACAGG |
| mTbp reverse | GTTGCTGAGATGTTGATTGCTG |
| mTubb3 forward | TAGACCCCAGCGGCAACTAT |
| mTubb3 reverse | GTTCCAGGTTCCAAGTCCACC |
| mNeuroD1 forward | ATGACCAAATCATACAGCGAGAG |
| mNeuroD1 reverse | TCTGCCTCGTGTTCCTCGT |
| mTet1 forward | CGGGTTTACAATGGCTCTTCG |
| mTet1 reverse | GGTTTGGGTGTGACTACTGGG |
| mTet3 forward | TGCGATTGTGTCGAACAAATAGT |
| mTet3 reverse | TCCATACCGATCCTCCATGAG |

**Supplementary Table 2 List of primary and secondary antibodies used for western blotting (WB) and immunofluorescence (IF) with the respective dilutions**

| Antibody | Distributor | Dilution |
|---|---|---|
| Anti-alphaTubulin | Sigma-Aldrich (T9025) | WB: 1:5000 |
| Anti-Bcat1 | Abcam (ab232700) | WB: 1:1000 |
| Anti-CD8a | R&D Systems (NBP2-52659) | IF: 1:100 |
| Anti-GFP | Abcam (ab13970) | IF: 1:500 |
| Anti-GFP | Biocat (AB011-EV) | IF: 1:500<br>WB: 1:1000 |
| Anti-Iba1 | FUJIFILM Wako Chemicals (019-1971) | IF: 1:250 |
| Anti-Ki67 | Abcam (ab15580) | IF: 1:250 |
| Anti-Sox2 | Merck Millipore (AB5603) | IF: 1:100 |
| Anti-TUBB31 | Biolegend (MMS-435) | IF: 1:100 |
| Anti-V5-tag | Cell Signaling Technology (13202S) | WB: 1:1000 |
| Anti-5-hmC | Active Motif (39770) | IF: 1:250 |
| Donkey anti-Chicken(H+L) CF488A | Sigma (SAB4600031-250ul) | IF: 1:1000 |
| Donkey anti-Chicken(H+L) CF633 | Sigma (SAB4600127-50ul) | IF: 1:1000 |
| Donkey anti-Goat(H+L) CF555 | Sigma (SAB4600059-250ul) | IF: 1:1000 |
| Donkey anti-Rabbit(H+L) CF647 | Sigma (SAB4600177-250ul) | IF: 1:1000 |
| Goat anti-Mouse (H+L) AF647 | ThermoFisher Scientific (A3272) | IF: 1:1000 |
| Goat anti-Mouse AF488 | ThermoFisher Scientific (A32723) | IF: 1:1000 |
| Goat anti-Rabbit AF555 | ThermoFisher Scientific (A21428) | IF: 1:1000 |
| HRP anti-Mouse | Cell Signaling Technology (7076S) | WB: 1:2500 |
| HRP anti-Rabbit | Cell Signaling Technology (7074S) | WB: 1:2500 |

**Supplementary Table 3 shRNA sequences used for pLKO.1-blast vector production and knockdown in mouse mGB2 cells**

| Oligo | Sequence |
|---|---|
| shTet1 1 forward | CCGGCAACTTGCATCCACGATTAATCTCGAGATTAATCGTGGATGCAAGTTGTTTTTG |
| shTet1 1 reverse | AATTCAAAAACAACTTGCATCCACGATTAATCTCGAGATTAATCGTGGATGCAAGTTG |
| shTet1 2 forward | CCGGTTTCAACTCCGACGTAAATATCTCGAGATATTTACGTCGGAGTTGAAATTTTTG |
| shTet1 2 reverse | AATTCAAAAATTTCAACTCCGACGTAAATATCTCGAGATATTTACGTCGGAGTTGAAA |
| shTet1 3 forward | CCGGCCTACGGGAAGCGACCATAATCTCGAGATTATGGTCGCTTCCCGTAGGTTTTTG |
| shTet1 3 reverse | AATTCAAAAACCTACGGGAAGCGACCATAATCTCGAGATTATGGTCGCTTCCCGTAGG |
| shTet3 1 forward | CCGGGCTCCAACGAGAAGCTATTTGCTCGAGCAAATAGCTTCTCGTTGGAGCTTTTTG |
| shTet3 1 reverse | AATTCAAAAAGCTCCAACGAGAAGCTATTTGCTCGAGCAAATAGCTTCTCGTTGGAGC |
| shTet3 2 forward | CCGGCTGATACCCTCCGGAAGTATGCTCGAGCATACTTCCGGAGGGTATCAGTTTTTG |
| shTet3 2 reverse | AATTCAAAAACTGATACCCTCCGGAAGTATGCTCGAGCATACTTCCGGAGGGTATCAG |
| shTet3 3 forward | CCGGGAACCTTCTCTTGCGCTATTTCTCGAGAAATAGCGCAAGAGAAGGTTCTTTTTG |
| shTet3 3 reverse | AATTCAAAAAGAACCTTCTCTTGCGCTATTTCTCGAGAAATAGCGCAAGAGAAGGTTC |
| shLuc forward | CCGGATGTTTACTACACTCGGATATCTCGAGATATCCGAGTGTAGTAAACATTTTTTG |
| shLuc reverse | AATTCAAAAAATGTTTACTACACTCGGATATCTCGAGATATCCGAGTGTAGTAAACAT |
